# Comparative small RNA and degradome sequencing provide new insights into antagonistic interactions in the biocontrol fungus *Clonostachys rosea*

**DOI:** 10.1101/2022.02.04.479213

**Authors:** Edoardo Piombo, Ramesh Raju Vetukuri, Poorva Sundararajan, Sandeep Kushwaha, Dan Funck Jensen, Magnus Karlsson, Mukesh Dubey

## Abstract

Necrotrophic mycoparasitism is an intricate process involving recognition, physical mycelial contact and killing of host fungi (mycohosts). During such interactions, mycoparasites undergo a complex developmental process involving massive regulatory changes of gene expression to produce a range of chemical compounds and proteins that contribute to the parasitism of the mycohosts. Small-RNAs (sRNAs) are vital components of post-transcriptional gene regulation, although their role in gene expression regulation during mycoparasitism remain understudied. Here, we investigated the role of sRNA-mediated gene regulation in mycoparasitism by performing sRNA and degradome tags sequencing of the mycoparasitic fungus *Clonostachys rosea* interacting with the plant pathogenic mycohosts *Botrytis cinerea* and *Fusarium graminearum* at two time points. The majority of differentially expressed sRNAs were down-regulated during the interactions with the mycohosts compared to a *C. rosea* self-interaction control, thus allowing de-suppression (up-regulation) of mycohost-responsive genes. Degradome analysis showed a positive correlation between high degradome counts and antisense sRNA mapping and led to the identification of 201 sRNA-mediated gene targets for 282 differentially expressed sRNAs. Analysis of sRNA gene targets revealed that the regulation of genes coding for membrane proteins was a common response against both mycohosts. While the regulation of genes involved in oxidative stress tolerance and cellular metabolic and biosynthetic processes was exclusive against *F*. *graminearum* highlighting common and mycohosts-specific gene regulation of *C*. *rosea*. By combining these results with transcriptome data collected in similar experimental conditions during a previous study, we expand the understanding of the role of sRNA in regulating interspecific fungal interactions and mycoparasitism.

**Importance:** Small-RNAs (sRNAs) are emerging as key players in pathogenic and symbiotic fungus-plant interactions, however, their role in fungal-fungal interactions remains elusive. In this study, we employed the necrotrophic mycoparasite *Clonostachys rosea* and plant pathogenic mycohots *Botrytis cinerea* and *Fusarium graminearum* and investigated the sRNA-mediated gene regulation in mycoparasitic interactions. The combined approach of sRNA and degradome tag sequencing identified 201 sRNA-mediated putative gene targets for 282 differentially expressed sRNAs highlighting the role of sRNA-mediated regulation of mycoparasitism in *C. rosea.* We also identified 36 known and 13 novel miRNAs and their potential gene targets at endogenous level, and at a cross-species level in *B. cinerea* and *F. graminearum* indicating a role of cross-species RNAi in mycoparasitism, representing a novel mechanism in biocontrol interactions. Furthermore, we showed that *C*. *rosea* adapts its transcriptional response, and thereby its interaction mechanisms, based on the interaction stages and identity of the mycohost.

## Introduction

RNA interference (RNAi) is a method of gene expression regulation based on small RNAs (sRNAs), which can influence gene regulation at the transcriptional and post-transcriptional level (1, 2). These sRNAs usually have a length of 18-40 nucleotides, and their silencing action is mediated mainly by three categories of enzymes: dicer or dicer-like endoribonucleases (DCLs), Argonaute (AGO) proteins and RNA-dependent RNA polymerases (RDRPs). The role of DCLs is to cleave double stranded RNA precursors, generating several categories of small RNAs, the most studied of which is micro RNAs (miRNAs, milRNAs in fungi). These are small non-coding RNAs of 18-26 nucleotides, normally generated from single stranded RNA forming hairpin structures (3).

These units are then recognized by the RNA induced silencing complex (RISC) and used as a guide by AGO proteins to cleave or inhibit the translation of transcripts showing complementarity to the sRNAs (4). As a final step, RDRPs generate additional double stranded RNAs from sRNAs, amplifying the silencing signal (5, 6). PhasiRNAs, extensively found in plants, are sRNAs of 21-26 nucleotides in size generated from the Dicer-driven cleavage of long precursors synthesized by RDRPs enzymes, acting on the cleaved targets of specific miRNAs (7, 8). sRNA-mediated transcript cleavage results in a rapidly degraded upstream fragment and a stable downstream fragment (9) and the resulting products of sRNA-directed transcript cleavage can be determined through the sequencing of the 5′-ends of uncapped polyadenylated mRNAs. PARE (parallel analysis of RNA ends) (10), also known as degradome sequencing, is a well-known technique to identify sRNA gene targets and cleavage sites by mapping the degraded reads on mRNA transcripts, and nucleotide base complementarity between the transcript and sRNA (Addo-Quaye et al., 2009) at 10^th^ and 11^th^ position is used to identify degradation peaks (11). Degradome sequencing combined with bioinformatic analysis has been used to identify candidate sRNAs and their putative gene target in plants (10, 12, 13) as well as in fungi such as *Fusarium graminearum* and *Rhizophagus irregularis* (14, 15).

RNA interference has been observed to influence multiple processes at the endogenous level in fungi, such as sexual reproduction in *F. graminearum* (14), response to cellulose in *Trichoderma reesei* (16), mycelial growth and conidiogenesis in *Metarhizium anisopliae* (17) and *T. atroviride* (18). In addition to their endogenous role, specific sRNAs can travel between interacting species and regulate the function of recipient cells by hijacking their RNAi machinery (19, 20). Furthermore, RNAi has recently been exploited to control plant pathogens by exogenous application of sRNAs targeting genes in pathogens essential for disease development by a process called spray-induced gene silencing (21–23) .

*Clonostachys rosea* is an ascomycete fungus known for its mycoparasitic and antagonistic ability against several plant pathogenic fungi (24) and nematodes (25). Therefore, some strains of *C. rosea* are commercialized and used for the biological control of plant diseases in crop production (24). The antagonistic ability of *C. rosea* is achieved through the production of specialized metabolites (26–29), hydrolytic enzymes (30–33) and other secreted proteins (34, 35). In addition, *C. rosea* possess numerous drug resistance membrane transporters that can mediate the expulsion of both endogenous and exogenous toxic compounds (36–40). During antagonistic interactions with other fungi, *C*. *rosea* can recognize its mycohosts and respond with both common and specific transcriptional changes (39), demonstrating a mycohost-dependent expression of the genetic machinery. However, the issue of how these changes in gene expression are mediated remained elusive until recent work demonstrated a role of DCL-mediated RNAi in antagonistic interactions in *C. rosea* (41).

The aim of this study was to i) expand the understanding of RNAi-mediated antagonistic interactions in *C. rosea* by identifying candidate sRNAs and their cleavage products (gene targets) at endogenous (within *C. rosea*) and cross-species (in mycohosts) level, and ii) to investigate if, and to what extent, *C*. *rosea* deploys common or mycohost-specific sRNAs in regulating antagonistic interactions. To achieve these objectives, we sequenced both sRNAs and degradome of *C. rosea* during two stages of interaction with two intrinsically and phylogenetically different plant pathogenic fungi (mycohosts), *Botrytis cinerea* and *F. graminearum*. By combining the results from sRNA and degradome sequencing with transcriptome data collected in similar experimental conditions during a previous study, we identify candidate sRNAs (including known and novel milRNAs) and their putative gene targets potentially associated with antagonistic interactions in *C. rosea*. This includes the identification of pathogen genes putatively cleaved by *C. rosea* sRNAs, already predicted in a previous study (41), indicative of cross-species transfer of sRNAs. Furthermore, comparative/combined sRNA and degradome analyses revealed that *C. rosea* could modulate its regulatory network depending on mycohosts and stages of non-self-interactions.

## Results

### Antagonistic effect of *C. rosea* against *B. cinerea* and *F. graminearum*

The antagonistic ability of *C*. *rosea* towards *B*. *cinerea* and *F. graminearum* was assessed by measuring the mycelial growth rate of interacting species in an *in vitro* dual culture-plate confrontation assay (Supplemental figure S1). The growth rate of each fungus grown alone (non-interaction) and against itself (self-interaction) were used as controls. In comparison to non-interaction control, no significant changes in mycelial growth rates of *C. rosea* or *B*. *cinerea* were found during self-interactions or non-self-interactions (Supplemental figure S2). In contrast, *F*. *graminearum* showed a significant (*P* ≤ 0.017) reduction in growth rate during non-self (CrFg) and self-interactions (FgFg) compared to the non-interaction (Fg) control three days post-inoculation (dpi). After four days of incubation, the mycelial growth rate of *F*. *graminearum* in non-self-interaction was reduced by 23 % (*P* = 0.010) compared to the non-interaction control (Supplemental figure S2). After 4 dpi, the mycelial fronts of *F*. *graminearum* during self-interaction merged, thereby preventing further measurements. The result is in line with the previous finding of Zapparata et al. (42), where lower growth rate of *F*. *graminearum* is also reported during self-interaction than when grown alone. After mycelial contact, *C. rosea* overgrew the mycelium of *B*. *cinerea* with the same rate as *C. rosea* non-interacting control. In contrast, there was a significant (*P* = 0.001) 53 % reduction in the *C. rosea* overgrowth rate on *F. graminearum* mycelium, compared to the growth rate in the non-interaction control or overgrowth on *B. cinerea* (Figure 1).

**Figure 1:**
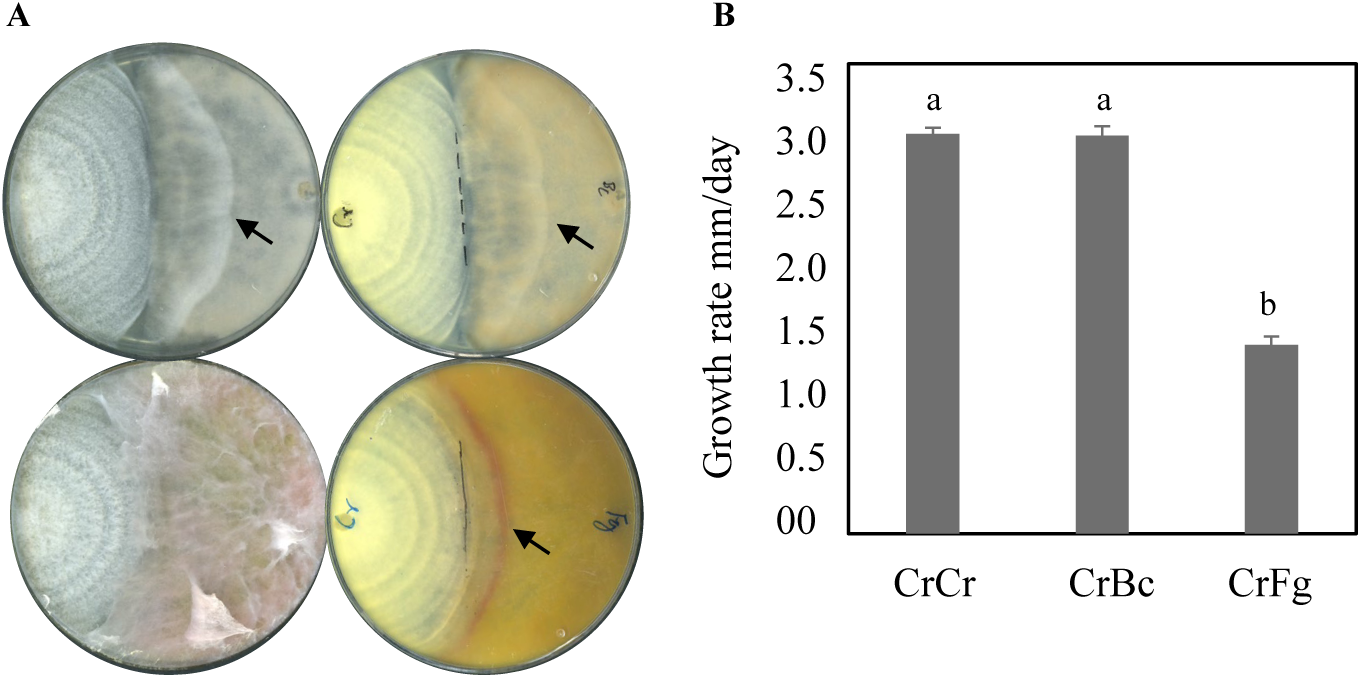
Measuring antagonistic ability of *C. rosea* against *B. cinerea* and *F*. *graminearum* using i*n vitro* dual culture plate confrontation assay. (**A**) Agar plug of *C*. *rosea* (right side in the plate) strains against *B. cinerea* or *F*. *graminearum* (left side in the plate) were inoculated on opposite sides in 9 cm diameter agar plates and incubated at 25°C. The experiment was performed in four replicates and photographs of representative plates were taken. An arrowhead indicates the *mycelial front of C. rosea*, solid and dashed line indicates the point of mycelial contact. (**B**) Growth rate (overgrow) of *C*. *rosea* WT and deletion strains on *B*. *cinerea* and *F*. *graminearum*. The growth of *C*. *rosea* on *B*. *cinerea* was measured from the point of mycelial contact. Error bars represent standard deviation based on four biological replicates. Different letters indicate statistically significant differences based on Tukey HSD method at the 95% significance level.

### Deep sequencing of *C. rosea* small RNAs

A dual culture-plate confrontation assay was used for total RNA extraction of *C*. *rosea* during *in vitro* interaction with two mycohosts, *B*. *cinerea* (CrBc) or *F*. *graminearum* (CrFg). Mycelial fronts were harvested at two stages, at mycelial contact and 24 hours after mycelial contact. *C*. *rosea* interacting with itself (CrCr) was used as a control treatment (Supplemental figure S1). sRNA sequencing generated a total of 1,052 million read pairs, ranging between 156 million to 192 million read pairs per treatment depending on the treatments (Supplemental table S1). After trimming of adaptor sequences, 969 million reads were obtained. The reads originating from *B*. *cinerea* or *F*. *graminearum* were removed by mapping sRNA reads to the *C*. *rosea*, *B*. *cinerea* and *F*. *graminearum* genomes, allowing zero mismatch. A total of 192 million read pairs from CrBc and CrFg unique to *C*. *rosea* (mapping to *C*. *rosea* genome and not to *B*. *cinerea* or *F*. *graminearum*) remained. Based on sRNA length that was previously observed for sRNAs in fungi (43), reads from 18 to 32 nucleotide length were used for further analyses.

### Origin and characteristics of *C. rosea* sRNAs

Analysis of sRNA length distribution showed no apparent differences in the proportion of size distribution of reads between the treatments. The 32 nt and 18 nt length represented the highest and lowest proportion of reads in all the treatments (Figure 2A). The analysis of the 5′ terminal nucleotide composition showed that a higher proportion of sRNA reads (42.6 %) starts with uracil (5’-U) during the contact stage of CrCr self-interaction compared with CrBc (39.0 %) and CrFg (39.1 %). Conversely, 24 h after contact the trend was opposite as a low proportion (39.5 %) of sRNAs with 5’-U was found during CrCr compared 42.7 % of reads with 5’-U during CrFg interactions (Figure 2B). To investigate the origin of sRNAs, we mapped sRNA sequences to the *C*. *rosea* genome. Our result showed that 70-76 % of sRNA sequences originated from exons (coding sequences [CDSs], 5’-untranslated regions [5’-UTRs] and 3’-UTRs), followed by tRNAs, promoter and intergenic regions (Figure 2C). A lower proportion (35 %) of sRNA reads was found to originate from CDSs 24 h after contact with *F. graminearum* (CrFg), compared with 44 % during CrCr self-interaction at the same stage. In addition, we analysed the relative proportion of sRNAs putatively originating from exons for their mapping to the sense or antisense strand of exons. Out of the total RNAs mapping on exons, 89 % and 11 % sRNA reads were mapped to the sense and antisense strand, respectively (Figure 2D). These results are in line with the previous findings obtained for *Mucor circinelloides* (44) and arbuscular mycorrhizal fungus *Rhizophagus irregularis* (15), *F. graminearum* (14) and *T. atroviride* (18) where sense strands of exons are shown to be the major source of sRNAs origin.

**Figure 2:**
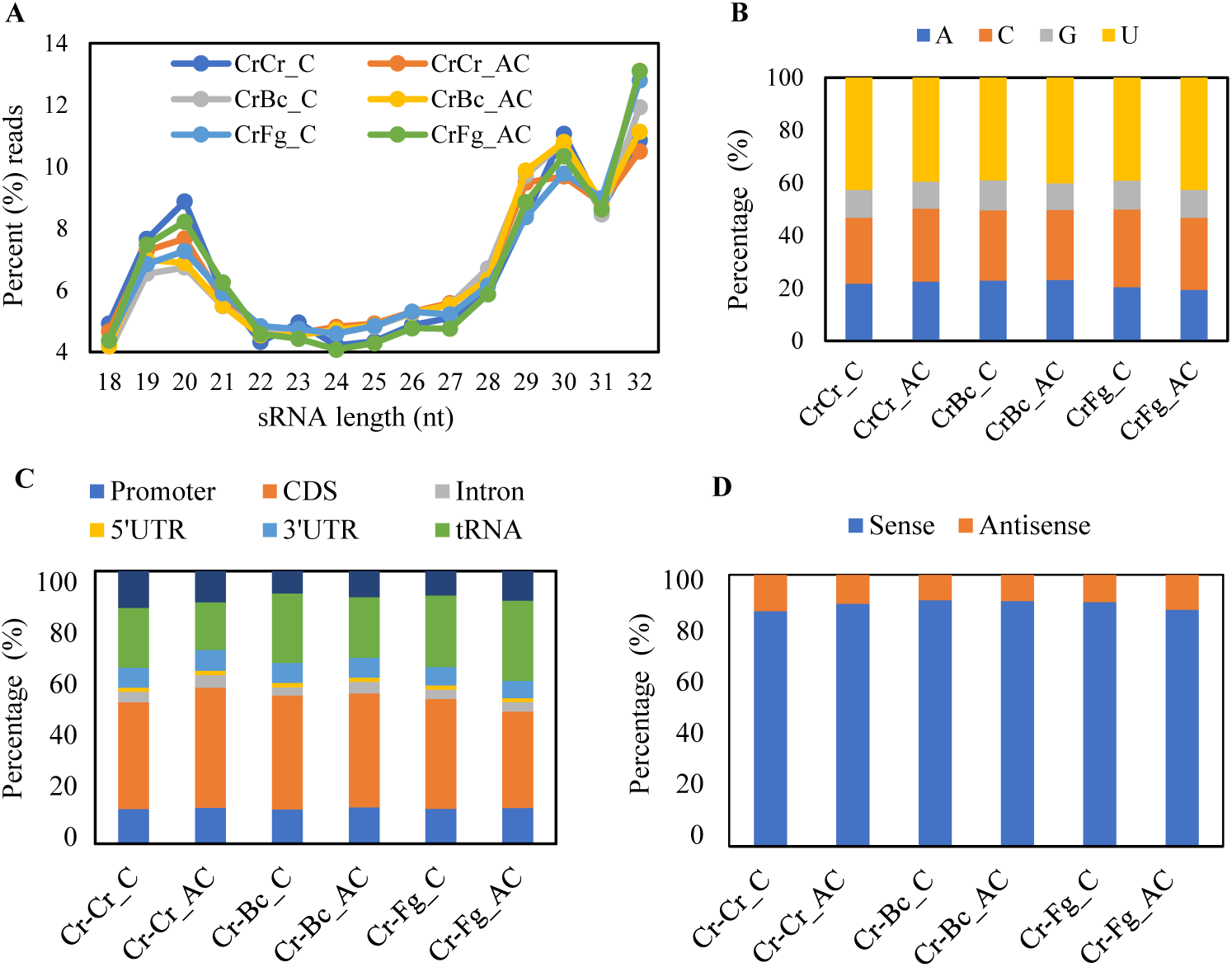
Characteristics of sRNAs in *C. rosea* interacting with *B. cinerea* (CrBc), F. graminearum (CrFg) and in the controls (CrCr), both at contact (C) and after contact (AC) stages of interaction. (**A**) length distribution, (**B**) 5’ nucleotide percentages, (**C**) location mapping, (**D**) sense mapping.

### sRNA expression in *C. rosea* is both mycohost and interaction stage dependent

To identify sRNAs differentially expressed in *C*. *rosea* during non-self-interactions, antisense-specific sRNAs and sRNAs potentially originating from intergenic and intronic regions were selected for differential expression analysis. Antisense-specific sRNAs were selected due to the reported high correlation between a high mapping of antisense sRNAs and high transcript degradation (14). On the other hand, sRNA originated from intergenic and intronic regions were selected due to previous findings where a majority of predicted *C. rosea* milRNAs were shown to locate in intergenic regions (41), and intron-containing mRNA precursors were shown to template siRNA synthesis in *Cryptococcus neoformans* (45). A summary of differentially expressed sRNAs is provided in Supplemental table S2.

The expression profile of sRNAs during CrBc and CrFg was compared to that of the CrCr control at the respective time points. In comparison to the CrCr control, 1947 and 564 sRNAs were down-regulated at contact and after contact stages of CrBc, respectively, while a lower number of sRNAs (590 and 36) were up-regulated under the same conditions. Among these, 269 down-regulated and 19 up-regulated sRNAs were common between both interaction stages (Figure 3). During the CrFg interactions, 2445 and 4250 sRNAs were down-regulated at contact and after contact stages, compared with CrCr control, while 790 and 257 were up-regulated under the same conditions. Among the sRNAs, 1232 were down-regulated at both time points, while 126 sRNAs were up-regulated at both time points (Figure 3). In summary, our data showed that differential sRNA expression in *C. rosea* is partially dependent on the developmental stage of the interaction, and the number of down-regulated sRNAs was higher than the number of up-regulated sRNAs.

**Figure 3:**
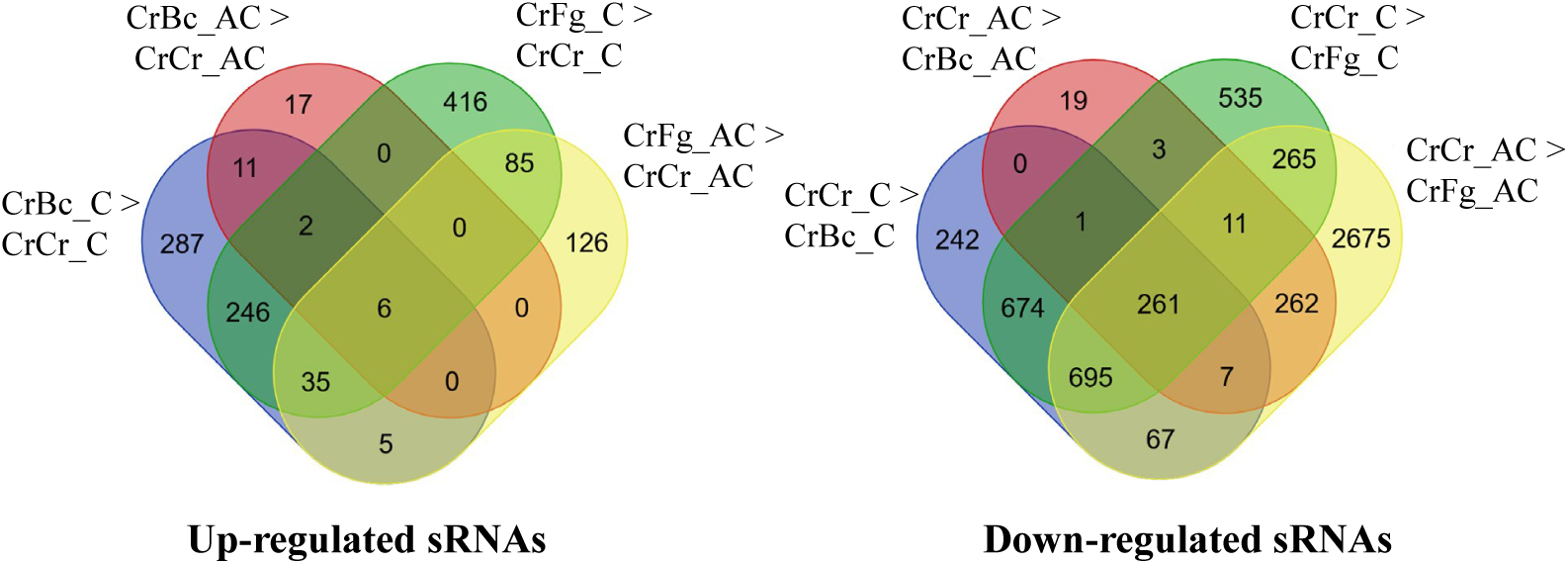
Venn diagram showing common and mycohost-specific expression of sRNAs during *C. rosea* interactions with *B. cinerea* (CrBc), *F. graminearum* (CrFg) compared to self-interaction (CrCr) control at contact (C) and after contact (AC) stage of interactions. (**A**) up-regulated sRNAs (**B**) down-regulated sRNAs.

Transcriptome (mRNA) analysis in *C*. *rosea* during CrBc and CrFg interactions showed common and species-specific responses towards the mycohosts (39). We investigated whether this finding held for sRNA expression and found that a higher number of down-regulated sRNAs (1631) were common at the contact stage against both mycohosts, whilst 316 and 814 sRNAs were specifically down-regulated against *B. cinerea* and *F. graminearum*, respectively.

After contact, 541 sRNAs were commonly down-regulated against both the mycohosts, while 23 and 3702 were uniquely expressed against *B. cinerea* and *F. graminearum*, respectively. Among the up-regulated sRNAs, 289 and 6 sRNAs were commonly up-regulated against both the mycohosts (Figure 3). In summary, sRNA expression analysis showed a higher number of *C. rosea* sRNAs were differentially expressed in CrFg compared to the number during CrBc. In addition, the number of differentially expressed sRNAs is higher at the contact stage in CrBc, but after contact in CrFg. Therefore, differential sRNA expression in *C. rosea* depends on both mycohost identity and interaction stage (Figure 3).

This result was further validated by a co-expression analysis executed with weighted correlation network analysis (WGCNA), which divided the differentially expressed sRNAs in 13 modules. The module eigengenes (ME), which are the first principal components of the expression matrix of each module, were shown to correlate to *B. cinerea* in the case of ME 6, 7 and 8, and to *F. graminearum* in the case of ME 3 and 4. ME 12 and 13 responded to the interaction stage, while ME 10 and 11 contained sRNAs up-regulated in the presence of any one of the two mycohosts (Figure 4A).

**Figure 4:**
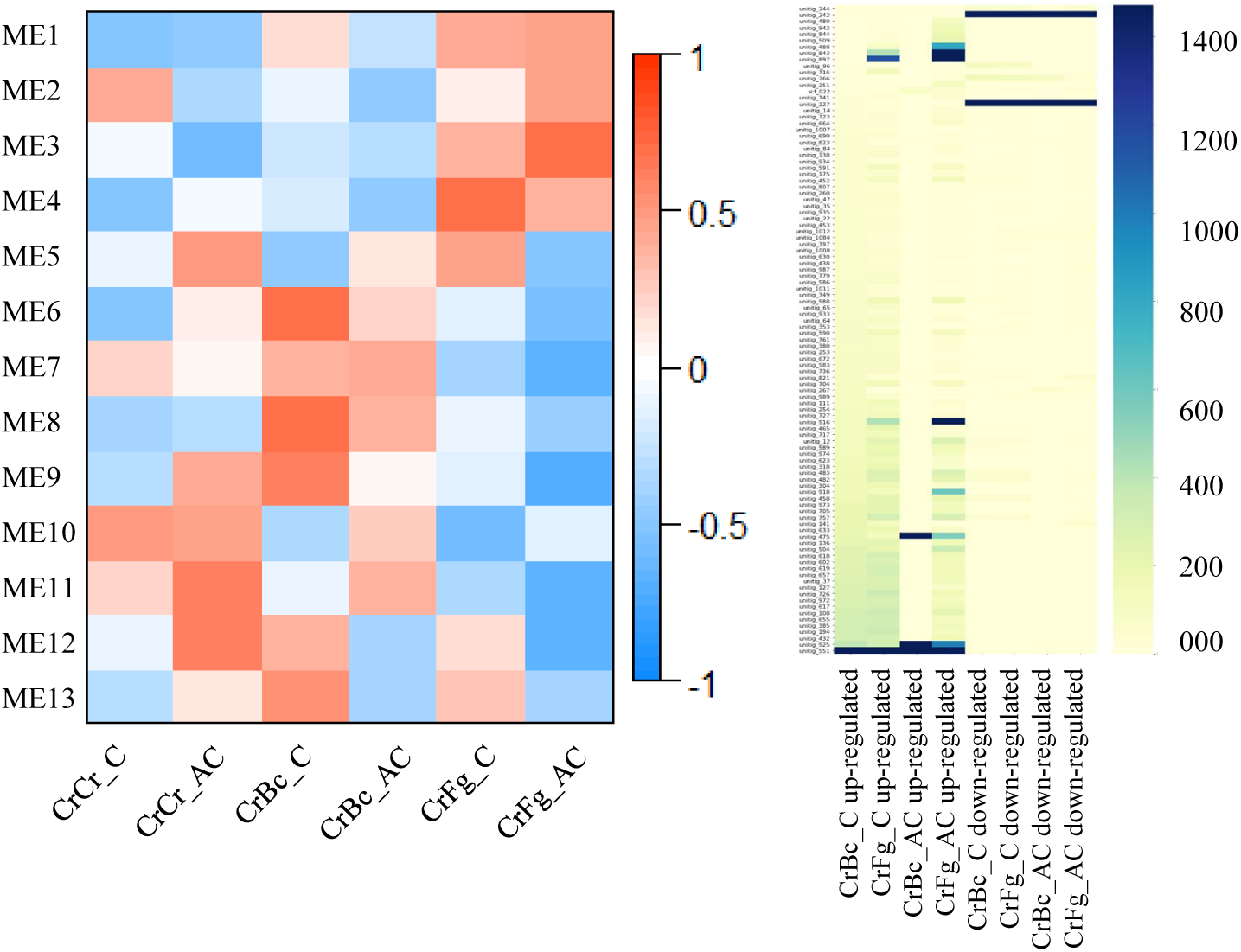
(**A**) The heatmap shows the spearman correlation between the module eigengenes of coexpression modules generated with WGCNA and the conditions examined in this study. *C. rosea* interaction with *B. cinerea* (CrBc), *F. graminearum* (CrFg) and the controls (CrCr), both at contact (C) and after contact (AC) stage of interactions. The modules were generated using the normalized expression values of differentially expressed sRNAs. (**B**) Heatmap showing the RPKM values obtained by mapping differentially expressed sRNAs to their scaffolds of origin. Only differentially represented scaffolds with a minimum RPM count of 70 are presented.

Differentially expressed sRNAs were mapped to the *C. rosea* genome to investigate whether these sRNAs originated from a specific region in the genome. sRNAs up-regulated at the contact stage of both mycohosts and after contact with *F. graminearum* mainly originated from a precise group of scaffolds (Figure 4B). The sRNAs up-regulated after contact with *B. cinerea* were too few to analyse. The down-regulated sRNAs had an even more specific origin, in most cases originating from either scaffold unitig_227 or unitig_242 (Figure 4B).

### Identification and expression analysis of *C. rosea* milRNAs

We used mirdeep2 (46) for milRNA prediction and identified 36 known and 13 novel milRNAs (Supplemental table S3). A summary of milRNAs sequences, origin, precursor and abundance is provided in Supplemental table S3. All 49 milRNAs had their reverse complement detected among the clean reads. Among those, 7 known and 4 novel milRNAs were differentially expressed in at least one of the examined situations (Table 1). In CrBc, five milRNAs (cro-mir-1, cro-mir-36, cro-mir-62, cro-mir-63 and cro-mir-70) were down-regulated while two milRNAs (cro-mir-4, cro-mir-72) were up-regulated. In CrFg, three milRNAs (cro-mir-1, cro-mir-36 and cro-mir-63) were down-regulated and five (cro-mir-8, cro-mir-9, cro-mir-23, cro-mir-34 and cro-mir-72) were up-regulated. Three (cro-mir-1, cro-mir-36 and cro-mir-63) of the down-regulated milRNAs were common to both CrBc and CrFg, while cro-mir-72 was up-regulated commonly to both interactions. The expression of a subset of sRNAs and miRNAs was further confirmed through stem-loop RT-qPCR (Table 2). A single milRNA (cro-mir-72) was upregulated to different degrees in the Cr-Bc C and Cr-Fg C data sets in both RNA-seq and stem-loop RT-qPCR analysis. The rest of the tested sRNAs and miRNAs were downregulated in both studies. In summary, the number of differentially expressed milRNAs at the contact stage of interaction (12 milRNAs) was higher than the number of differentially expressed milRNAs at after the contact stage (6 milRNAs, Table 1). Among the differentially expressed milRNAs, four (cro-mir-1, cro-mir-4, cro-mir-9, cro-mir-36) were proven to be DCL2-dependent as reported in a previous study of Piombo et al. (41). Their expression and predicted gene targets are presented in Table 3.

**Table 1:**
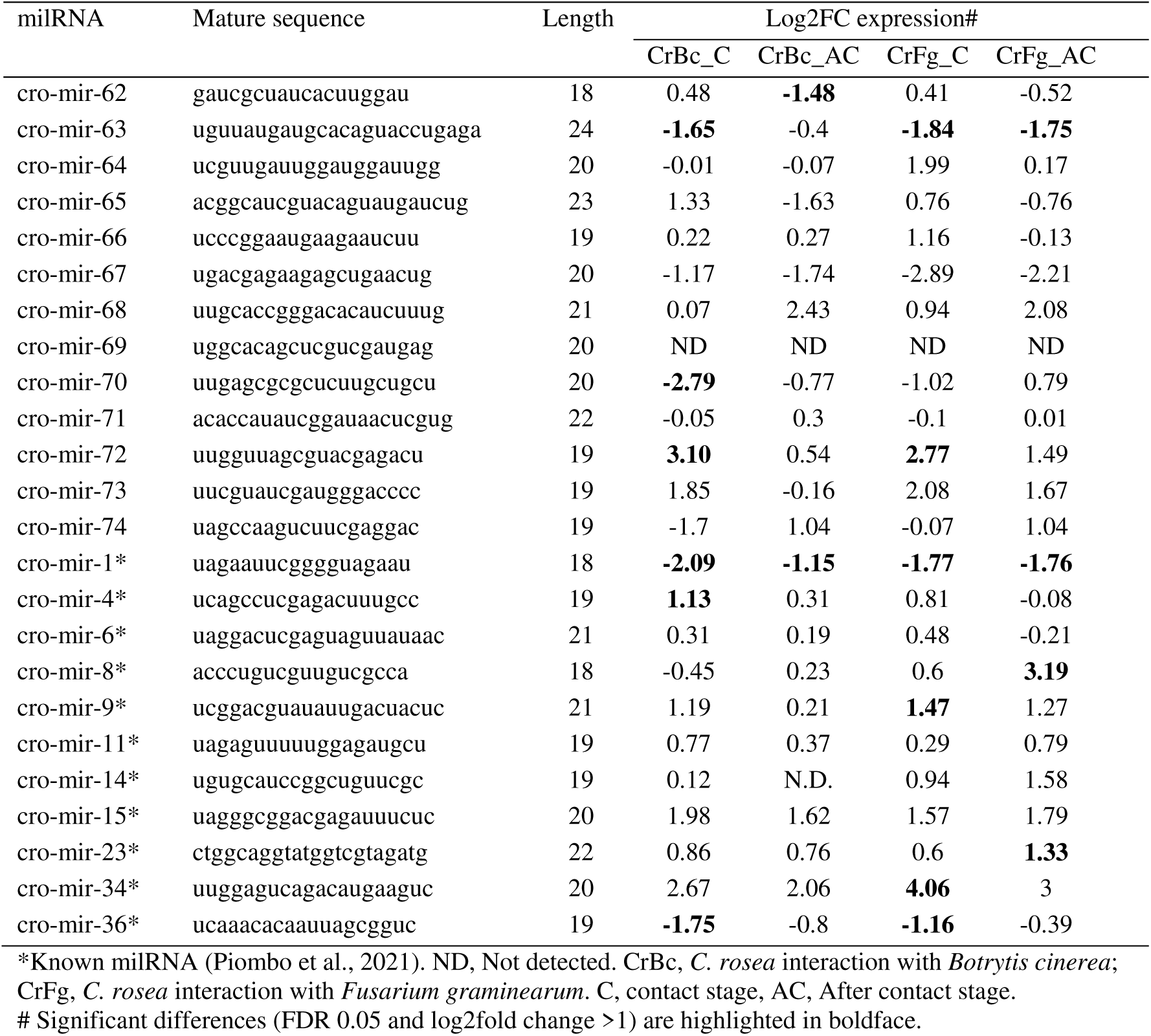
Novel and known detected milRNAs and their expression pattern in *C. rosea* during non-self-interactions with *B. cinerea* or *F. graminearium* compared to self-interaction control.

**Table 2:** Validation of sRNA sequencing through stem-loop RT-qPCR. A subset of sRNAs and miRNAs were selected for the validation.

| sRNAs ID | Log2FC Sequencing |  | Log2FC stem-loop RT-qPCR |  |
| --- | --- | --- | --- | --- |
|  | Cr-Bc C | Cr-Fg C | Cr-Bc C | Cr-Fg C |
| ae_seq_277875_x17263 | -4.06 | -3.4 | -4.02 | -3.92 |
| ii_seq_8561252_x801 | -3.96 | -4.03 | -2.82 | -5.29 |
| cro-mir-1 | -2.08 | -1.77 | -3.98 | -2.27 |
| cro-mir-36 | -1.75 | -1.16 | -1.82 | -3.04 |
| cro-mir-72 | 3.1 | 2.77 | 0.56 | 0.48 |

CrBc, *C. rosea* interaction with *Botrytis cinerea*; CrFg, *C. rosea* interaction with *Fusarium graminearum*. C, contact stage, AC, After contact stage.

**Table 3:** Differential expression of DCL2-dependent milRNAs, and their gene targets.

| milRNAs# | Log2FC sRNA expression* |  |  |  | Putative gene targets | Annotation |
| --- | --- | --- | --- | --- | --- | --- |
|  | CrBc_C | CrBc_AC | CrFg_C | CrFg_AC |  |  |
| Endogenous gene targets |  |  |  |  |  |  |
| cro-mir-1 | -2.08 | -1.14 | -1.77 | -1.76 | CRV2T00003756 | aminoacyl-tRNA ligase |
| cro-mir-1 | -2.08 | -1.14 | -1.77 | -1.76 | CRV2T00017618 | Uncharacterized |
| cro-mir-36 | -1.75 | -0.8 | -1.16 | -0.39 | CRV2T00011823 | sulfuric ester hydrolase |
| cro-mir-36 | -1.75 | -0.8 | -1.16 | -0.39 | CRV2T00013380 | ATPase |
| cro-mir-36 | -1.75 | -0.8 | -1.16 | -0.39 | CRV2T00005499 | Uncharacterized |
| cro-mir-36 | -1.75 | -0.8 | -1.16 | -0.39 | CRV2T00000111 | Uncharacterized |
| cro-mir-36 | -1.75 | -0.8 | -1.16 | -0.39 | CRV2T00014914 | Uncharacterized |
| cro-mir-36 | -1.75 | -0.8 | -1.16 | -0.39 | CRV2T00000903 | Uncharacterized |
| cro-mir-36 | -1.75 | -0.8 | -1.16 | -0.39 | CRV2T00004261 | Uncharacterized |
| cro-mir-4 | 1.13 | 0.31 | 0.81 | -0.08 | CRV2T00011242 | Uncharacterized |
| cro-mir-9 | 1.19 | 0.21 | 1.47 | 1.27 | CRV2T00004339 | SNF2 family helicase |
| Cross-species gene targets in *B. cinerea* |  |  |  |  |  |  |
| cro-mir-4 | 1.13 | 0.31 | 0.81 | -0.08 | XM_024690414.1 | Chitin synthase<br>BcCHSIV |
| cro-mir-4 | 1.13 | 0.31 | 0.81 | -0.08 | XM_024693385.1 | Serine threonine-protein<br>phosphatase |
| cro-mir-4 | 1.13 | 0.31 | 0.81 | -0.08 | XM_024696641.1 | Vacuolar sorting protein<br>Bcvps27 |
| cro-mir-4 | 1.13 | 0.31 | 0.81 | -0.08 | XM_024695521.1 | Chromatin binding factor<br>Bcyta7 |
| cro-mir-4 | 1.13 | 0.31 | 0.81 | -0.08 | XM_001556587.2 | Transcription factors<br>bcltf3 |
| Cross-species gene targets in *F. graminearum* |  |  |  |  |  |  |
| cro-mir-9 | 1.19 | 0.21 | 1.47 | 1.27 | XM_011320613.1 | Zinc-binding protein |
| cro-mir-9 | 1.19 | 0.21 | 1.47 | 1.27 | XM_011329717.1 | sec1 family vesicle<br>transport protein |

Cr, *C. rosea*; Bc, *B. cinerea*; Fg, *F. graminearum*; C, contact stage; AC, After contact stage.
\*Significant differences (FDR 0.05 and log2fold change > 1) are highlighted in boldface.

### Degradome analysis showed a positive correlation between high degradome counts and antisense sRNA mapping

The *C. rosea* degradome samples were sequenced to identify sRNA gene targets, 5’ non-capped degradation products of mRNA, producing between 4.2 and 10.7 million clean reads depending on the sample (Supplemental table S4). A high proportion of the reads (62 % on average) from *C. rosea* and the mycohosts’ interaction was mapped to *C. rosea*. In comparison, the average count for *B. cinerea* and *F. graminearum* were 31 % and 32 %, respectively. Curiously, the number of reads assigned to *F. graminearum* was higher in samples at the 24 h after contact time point (34 %) compared with the contact stage (29 %). The opposite tendency was observed in degradome reads from CrBc where the higher proportion of reads (38 %) mapped to *B. cinerea* at the contact stage compared with 25 % after contact (Supplemental table S4). Mapping of degradome reads to the *C. rosea* genome showed that the majority of reads were mapped on the transcribed regions of the genome, with 54 % aligning to CDS regions, 16 % to 3’-UTRs and 1.6 % to 5’-UTRs. Moreover, 4.9 % of the reads mapped to promoter sequences, while 13.6 % was assigned to intergenic regions. A meager fraction of sequences (0.01 %) was successfully mapped to known tRNAs (Figure 5A). Degradome-based hierarchical clustering grouped degradomes from the CrBc and CrFg contact stage together. In contrast, a higher similarity between the degradome from CrBc after contact stage and self-interaction (CrCr) was found (Figure 5B).

**Figure 5:**
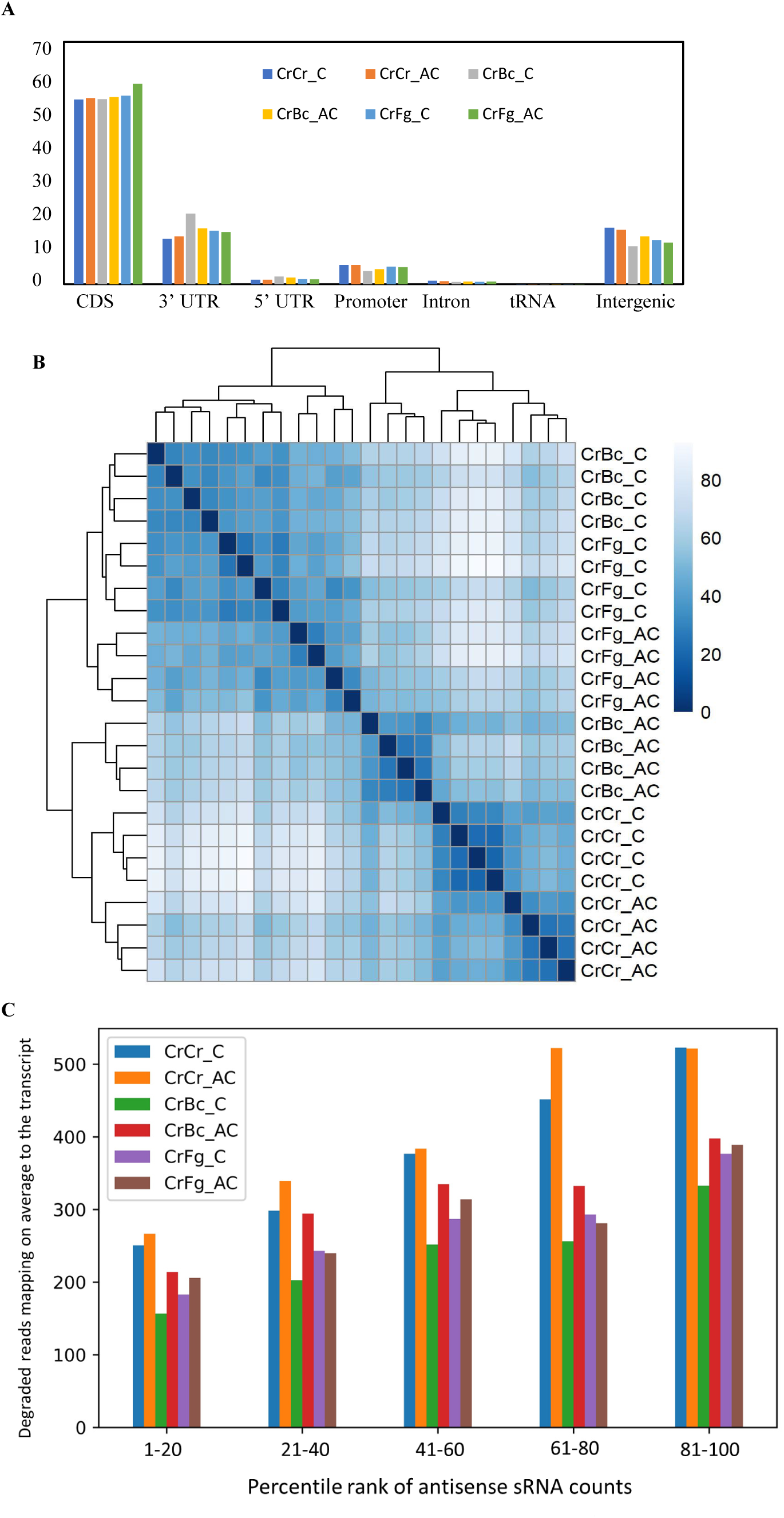
Degradome sequence analyses. **(A)** results of degradome read mapping on gene features. The reads belong to *C. rosea* interacting with *B. cinerea* (CrBc), *F. graminearum* (CrFg) and controls (CrCr). (**B)** Hierarchical clustering of samples depending on degradome read mapping to *C. rosea* transcripts. (**C**) Average degraded read count of *C. rosea* genes, depending on their percentile rank of antisense sRNA counts. Percentile ranks were assigned to each gene based on its antisense sRNA counts. Genes with an antisense sRNA count of zero were not considered. The reads belong to *C. rosea* interacting with *B. cinerea* (CrBc), *F. graminearum* (CrFg) and in the controls (CrCr), C, contact, AC after contact stage of interaction.

We analysed the correlation between sRNAs with antisense orientation mapping and transcript cleavage to analyse sRNA-mediated transcript degradation. The genes were divided into 4 groups depending on their antisense sRNA counts, with each group comprising 25 % of all genes and containing genes with a sRNA antisense count higher than the genes in the previous group. Then, the average degradome count was observed for each group in each interaction. Among the 25 % of genes with lower antisense sRNA counts, the average degradome read count was between 160 and 270, while the same value was between 300 and 500 for the genes in the last group. This finding showed a positive correlation between mapping of antisense sRNAs to the transcripts and higher degradome count (Figure 5C) and corroborated the use of degradome sequencing for investigating sRNA-mediated gene regulation.

### Identification of endogenous and cross-species gene targets using degradome sequencing

Transcriptome-wide degradome analysis has previously been used for large-scale sRNAs target identification (13, 14). We used CleaveLand (11) on the degradome data for detection of transcripts with a higher-than-average degradome count at the point of alignment with a differentially expressed sRNA (FDR < 0.05 and log2(FC) ≥ 1). Later, it was also confirmed through evaluation of differential degradation of the targets (FDR < 0.05 and log2(FC) ≥ 1) at the predicted point of alignment between sRNAs and transcripts. In total we identified 201 putatively cleaved endogenous genes for 282 differentially expressed sRNAs (Table 4; Supplemental table S5). Target plots showing comparative transcript cleaving of 10 sRNAs gene targets are presented in Figure 6.

**Figure 6:**
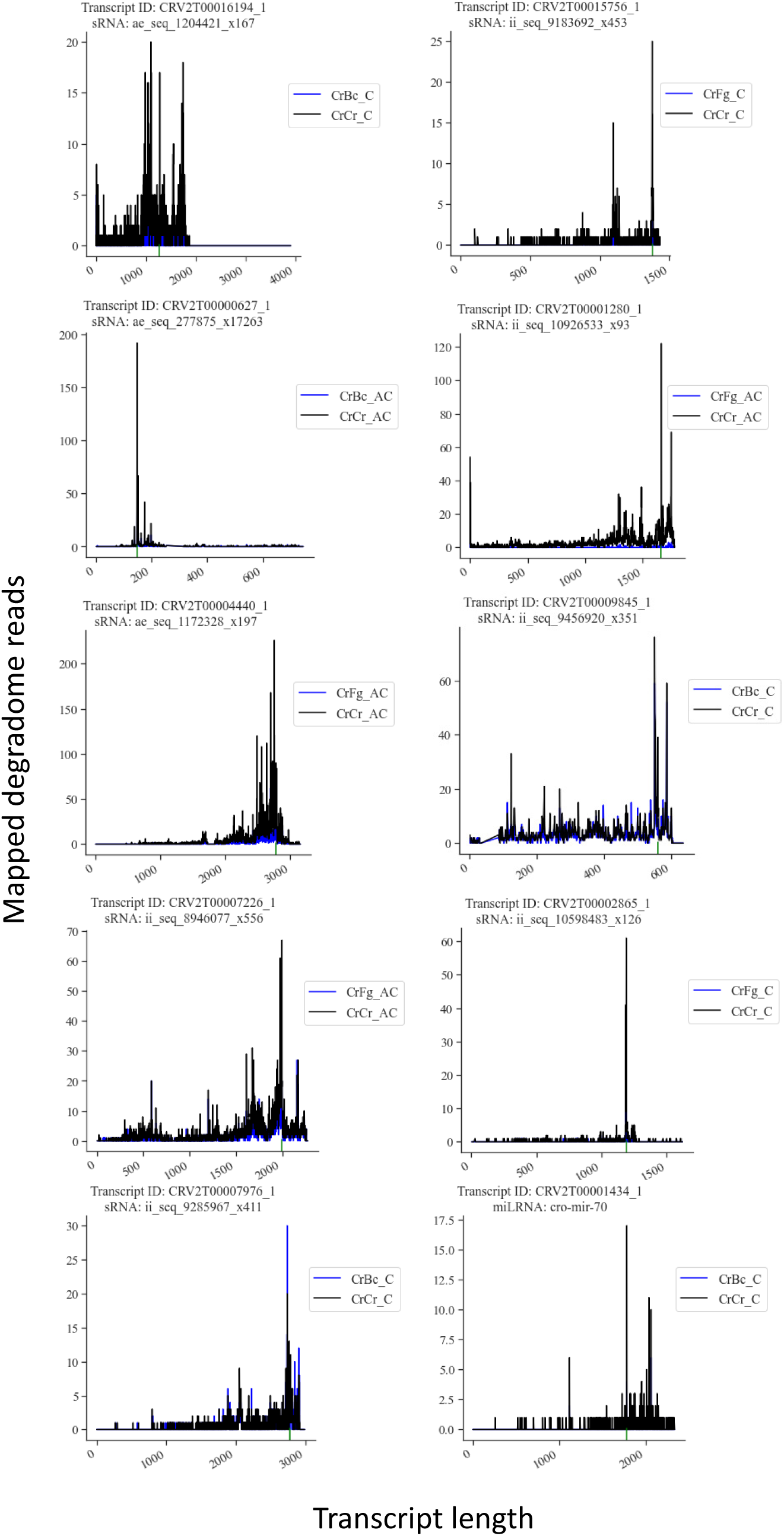
Target plots showing sRNA-mediated transcript cleavage. The mapping of degradome reads to gene of interest are shown. Green lines under the X axis indicate the area of alignment between the transcript and the considered sRNA. Transcript ID of gene and sRNA identification number is shown for each plot.

**Table 4:** Differentially expressed sRNAs in *C. rosea* during non-self-interactions with *B. cinerea* or *F. graminearium* compared to self-interaction control, and their endogenous gene targets of interest.

| sRNA identifier | Log2FC sRNA expression#* |  |  |  | Putative gene target | Gene target annotation |
| --- | --- | --- | --- | --- | --- | --- |
|  | CrBc_C | CrBc_AC | CrFg_C | CrFg_AC |  |  |
| Endogenous gene targets |  |  |  |  |  |  |
| ae_seq_1204421_x167 | -1.83 | -0.06 | -1.24 | -0.39 | CRV2T00016194 | MFS transporter |
| ii_seq_9183692_x453 | -1.23 | -0.55 | -1.62 | -0.98 | CRV2T00015756 | Catalase isozyme P |
| ae_seq_277875_x17263 | -4.06 | -1.67 | -3.4 | -3.58 | CRV2T00000627 | Autophagy 5 homolog |
| ii_seq_10926533_x93 | -3.7 | -2.01 | -2.8 | -2.87 | CRV2T00001280 | Chitinase GH18 |
| ae_seq_1172328_x197 | 0.07 | 0.06 | 0.21 | -2.18 | CRV2T00004440 | MAP kinase |
| ii_seq_9456920_x351 | -1.94 | -0.24 | -0.48 | -1.49 | CRV2T00009845 | Transcription factor |
| ii_seq_8946077_x556 | -3.56 | -1.14 | -3.1 | -2.43 | CRV2T00007226 | Transcription factor |
| ii_seq_10598483_x126 | -1.85 | -0.28 | -2.69 | -1.96 | CRV2T00002865 | Transcription initiation factor |
| ii_seq_9285967_x411 | 2.74 | 0.25 | 2.8 | 0.27 | CRV2T00007976 | Involved in glycogen biosynthesis |
| cro-mir-70 | -2.79 | 0.77 | -1.02 | 0.79 | CRV2T00001434 | Membrane transporter |
| Cross-species gene targets in *F. graminearum* |  |  |  |  |  |  |
| ii_seq_9854567_x248 | -0.12 | 0.48 | 2.38 | 3.35 | XM_011326499.1 | Rab gdp-dissociation inhibitor |
| cro-mir-72 | 3.1 | 0.54 | 2.77 | 1.49 | XM_011323146.1 | Elongation factor 3 |

Cr, *C. rosea*; Bc, *B. cinerea*; Fg, *F. graminearum*; C, contact stage; AC, After contact stage.

We identified 47 and 13 gene targets putatively cleaved by 64 and 16 down-regulated sRNAs at contact and after contact time points, respectively, during CrBc (Figure 7A). Among these, three gene targets were common between the two time points. Seventeen transcripts, targeted by 21 up-regulated sRNAs, were predicted in CrBc at contact, while no targets were predicted for the sRNA up-regulated after contact (Figure 7B). Compared to CrBc, the number of targets detected during the interaction with *F. graminearum* was higher. We found 94 gene targets for 111 down-regulated sRNAs at the contact stage, and 103 putative cleaved transcripts targeted by 163 down-regulated sRNAs at after contact stage (Figure 7A). Thirty-eight gene targets were common between the two time points. For sRNAs up-regulated during CrFg, we identified 18 cleaved transcripts targeted by 21 sRNAs at the contact stage, while the number of putative gene targets dropped to four at the after contact time point, predicted to be cleaved by three up-regulated sRNAs (Figure 7B). In summary, analyses of degradome data corroborate the mycohost and interaction stage-dependent response of *C*. *rosea* during non-self-interactions.

**Figure 7:**
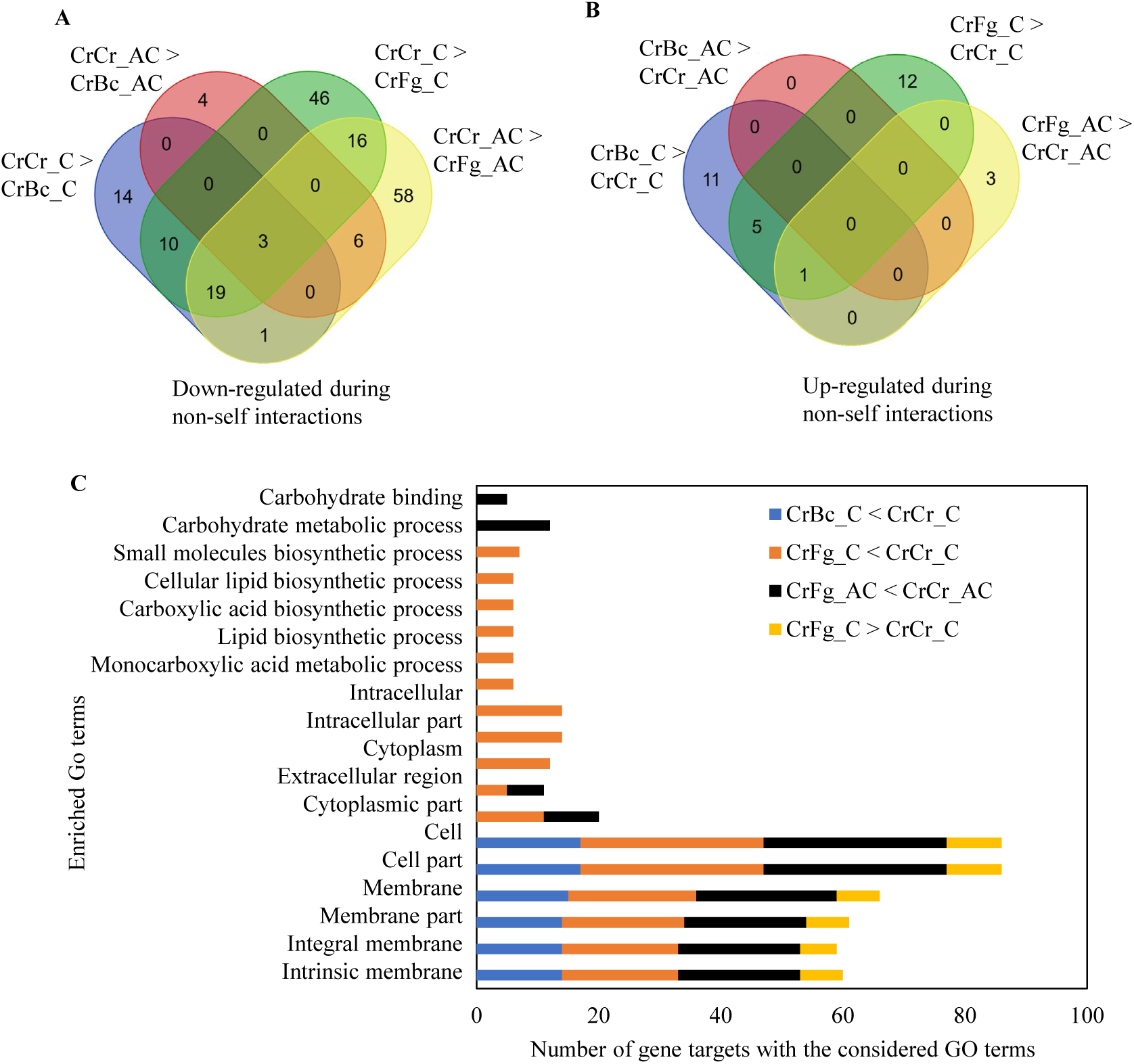
Distribution of *C. rosea* transcripts putatively cleaved by sRNA up-regulated (**A**) and down-regulated (**B**) during *C. rosea* interaction with *B. cinerea* (CrBc), *F. graminearum* (CrFg) compared with self-interaction controls (CrCr), both at contact (C) and after contact (AC) stage of interactions. The cleavage by up-regulated or down-regulated sRNAs is confirmed by corresponding over degradation or under degradation at the alignment point between the transcript and the sRNA. (**C**) Distribution of GO terms enriched in *C. rosea* transcripts putatively cleaved by sRNA down-regulated during interactions with *B. cinerea* or *F. graminearum*.

Additionally, we used degradome data from the interacting mycohosts and searched for potential gene targets of differentially expressed *C. rosea* sRNAs in *B. cinerea* and *F*. *graminearum*. Our result identified 43 and 91 potential gene targets in *B. cinerea* and *F. graminearum* with higher-than-average degradome read count in the mapping site of a differentially expressed *C. rosea* sRNA (Supplemental table S6). The 43 *B. cinerea* transcripts were putatively cleaved by 40 sRNAs up-regulated at contact stage. Among *F. graminearum* gene targets, 65 gene targets were putatively targeted by 70 sRNA up-regulated at contact stage, 13 by 20 sRNA up-regulated only after contact and 13 by 6 sRNAs up-regulated at both time points.

### Gene ontology enrichment analyses

sRNA gene targets were used for gene ontology (GO) term analysis to investigate biological processes, cellular components and molecular functions enriched among sRNA gene targets during the interspecific interactions in *C. rosea*. During CrBc, GO terms GO:0016020 (membrane), GO:0016021 (integral to membrane) and GO:0031224 (intrinsic to membrane) were enriched among the targets of sRNAs down-regulated at contact stage of interaction, suggesting a role of sRNA-mediated gene silencing in regulating expression of membrane proteins, such as the major facilitator superfamily (MFS) transporter CRV2T00016194 (Figure 7C, Table 4). No GO terms were enriched among transcripts putatively cleaved during the after contact stage of CrBc. Similarly, no GO terms were enriched during CrBc among the targets of up-regulated sRNAs, at any time point. However, the transcript CRV2T00007976, a glycoside hydrolase involved in glycogen biosynthesis, was predicted to be cleaved by ii_seq_9285967_x411 up-regulated at the contact stage (Figure 7C, Table 4).

During CrFg, the terms GO:0032787 (monocarboxylic acid metabolic process), GO:0008610 (lipid biosynthetic process), GO:0016053 (organic acid biosynthetic process), GO:0046394 (carboxylic acid biosynthetic process), GO:0044255 (cellular lipid metabolic process) and GO:0044283 (small molecule biosynthetic process) were enriched among targets of sRNAs down-regulated at contact in CrFg (Figure 7C). Since transcripts cleavage is a negative form of regulation, this points to an up-regulation of primary metabolism in *C. rosea* upon contact with *F. graminearum*. In CrFg 24 h after contact, the enriched GO terms among targets of down-regulated sRNAs were GO:0005975 (carbohydrate metabolic process) and GO:0030246 (carbohydrate binding). Regarding cellular localization, the terms GO:0005576 (extracellular region), GO:0016020 (membrane), GO:0016021 (integral to membrane) and GO:0031224 (intrinsic to membrane) were enriched among the putatively cleaved targets in CrFg both at contact and after contact (Figure 7C), suggesting that *C. rosea* responds to *F. graminearum* by reducing the cleaving of transcripts that encode for proteins able to interact directly with the mycohost through secretion or membrane localization. Response to oxidative stress was also predicted to be involved in response to *F. graminearum*, as the catalase isozyme P (CRV2T00015756) was among the transcripts putatively cleaved by sRNAs (ii_seq_9183692_x453) down-regulated in CrFg at contact (Table 4, Figure 7C). Among the targets of up-regulated sRNAs, the GO terms GO:0016020 (membrane), GO:0016021 (integral to membrane) and GO:0031224 (intrinsic to membrane) were again enriched during CrFg at contact, while no GO term was predicted at the after contact stage (Figure 7C). This suggests that the response to *F. graminearum* in *C. rosea* involves a rapid shift of membrane proteins to mediate the interaction with the mycohost.

### Identification of endogenous and cross-species gene targets of *C. rosea* milRNAs

*C. rosea* down-regulated milRNAs were predicted to cleave seven endogenous genes for four down-regulated milRNAs (Supplemental table S7). Among these, we found genes targets putatively coding for an aminoacyl-tRNA ligase (CRV2T00003756_1), a sulfuric ester hydrolase (CRV2T00011823_1) and an ATPase (CRV2T00013380_1). Up-regulated milRNAs, on the other hand, were predicted to cleave one endogenous transcript (CRV2T00001434) encoding a putative transmembrane protein (Supplemental table S7).

Cross-species gene targets analysis showed that cro-mir-72 was predicted to cleave the *F. graminearum* transcript XM_011323146.1 coding for elongation factor 3 (Table 4, Supplemental table S8). The role of elongation factor 3 in oxidative resistance is characterized in *Saccharomyces cerevisiae* (47).

### Mycohost-responsive milRNAs are not well conserved in *Clonostachys* spp.

To evaluate milRNA conservation in the genus *Clonostachys*, subgenus *Bionectria*, we searched for milRNA precursor sequences in genomes of five other species sequenced until date (48). Among the 74 milRNAs observed in this study and in that of (41), nine milRNAs were detected in at least five of the six *Clonostachys* spp, while 44 were detected in at least one more *Clonostachys* sp. (Supplemental table S9; Figure 8A). However, the differentially expressed milRNAs seemed to be less conserved than average (Figure 8B), with almost half of them detected only in *C. rosea*, while the others tended to be also observed in the more closely related *C. chloroleuca, C. byssicola* or *C. rhizophaga*.

**Figure 8.**
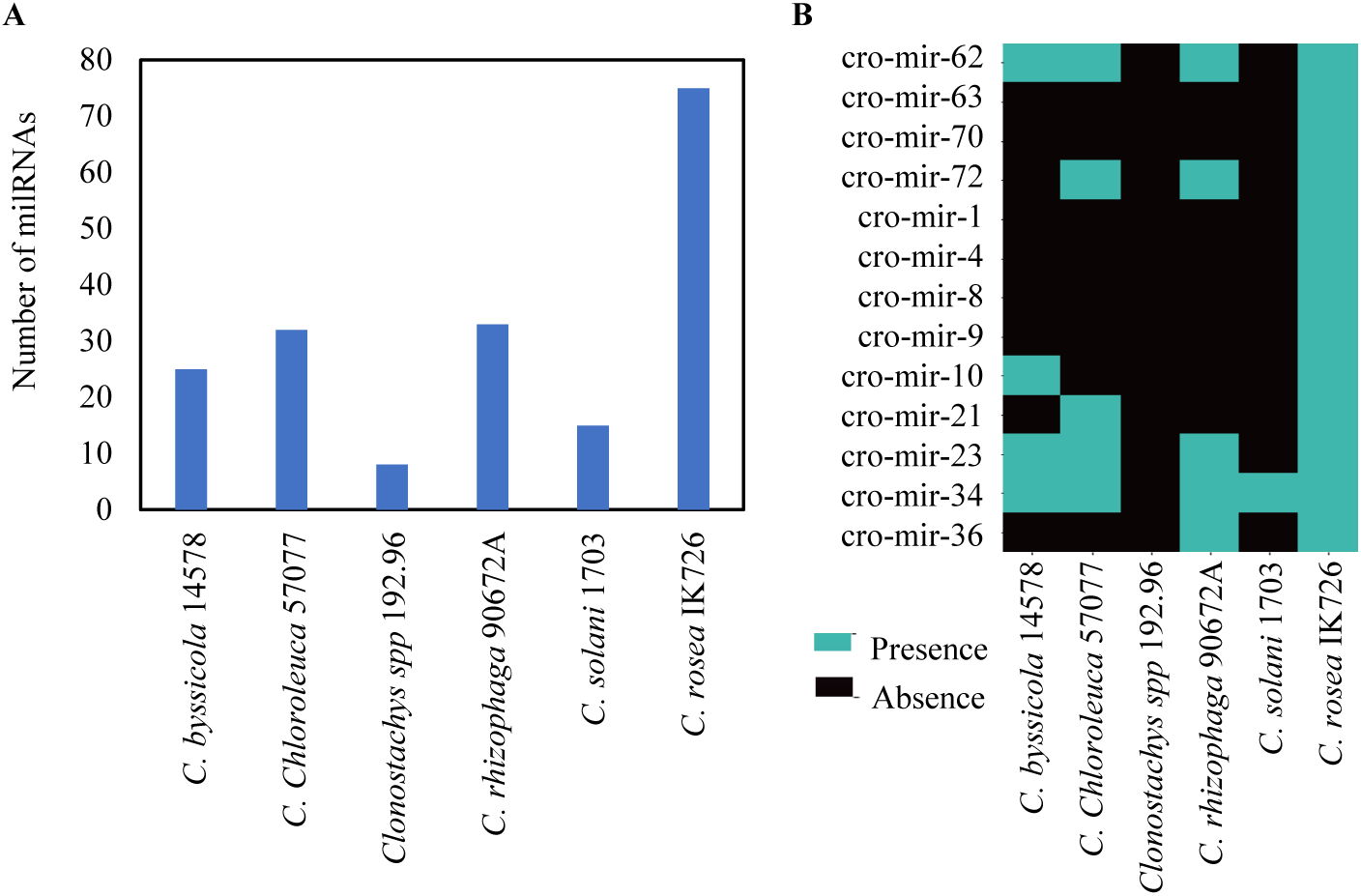
milRNA distribution in *Clonostachys* spp. (**A**) The number of *C. rosea* milRNAs detected in other *Clonostachys* spp. (**B**) Distribution of differentially produced *C. rosea* milRNAs during interactions with *B. cinerea* or *F. graminearum* compared with self-interactions control (CrCr).

### Validation of degradome-based gene targets by transcriptome sequencing

We used transcriptome data from our previous work collected at the contact stage of CrBc and CrFg interactions (41). We analysed the expression pattern of putative gene targets identified in this study for validation. In Piombo et al. (41), transcriptome (mRNAs and sRNAs) of *C*. *rosea* wild type (WT), *dcl1* deletion mutant (Δ*dcl1*) and *dcl2* deletion mutant (Δ*dcl2*) were sequenced during *in vitro* interactions with *B. cinerea* or *F. graminearum*. We expected that sRNA-cleaved transcripts identified from degradome sequencing to be de-suppressed (consequently up-regulated) in the Δ*dcl1* and Δ*dcl2* strains, compared to the WT, lacking sRNAs required for transcript cleavage. This approach was able to verify up to 37 % of the degradome-based gene targets, depending on the examined condition. Our result showed that a higher proportion of gene targets were verified for down-regulated sRNAs (Supplemental figure S3) compared to up-regulated sRNA gene targets. This positively correlates with sRNAs expression pattern and degradome-based target prediction, as a higher proportion of sRNAs were down-regulated during non-self-interactions and, consequently, a higher number of degradome-based gene targets were predicted for down-regulated sRNAs. These results suggest that the degradome-sequencing can be used for sRNA gene target identification.

### Identification of phasiRNAs

PHASIS analysis identified 46 phasiRNAs in *C. rosea*, belonging to 7 families (Supplemental table S10) originating mainly from tRNAs. Although 35 phasiRNAs were differentially expressed in *C. rosea* in at least one stage of interspecific interactions, compared with the CrCr self-interactions, no alignment between these sequences and the degradome data was found.

## Discussion

*Clonostachys rosea* is a necrotrophic mycoparasite with broad range of mycohosts (24). A transcriptome study of *C. rosea* during interactions with *B. cinera* and *F. graminearum* showed both common and specific response (39). The difference in transcriptomic response is considered to be associated with the intrinsic differences of the mycohosts, for instance a differential composition of the cell wall and the ability to produce a different spectrum of specialized metabolites, hydrolytic enzymes, reactive oxygen species and other virulence factors. This was confirmed by a study of Piombo et al. (41), which showed that mycohosts *B. cinerea* and *F. graminearum* responded differently against *C. rosea* on a transcriptional level. Genes involved in synthesizing and transporting specialized metabolites, including the polyketide fusarielin, and mycotoxins like zearalenone and deoxynivalenol (DON), were up-regulated in *F. graminearum* during interaction with *C. rosea* WT compared to a *C. rosea* Δ*dcl2* mutant strain (41). In *B*. *cinerea*, genes involved in cell wall biosynthesis were up-regulated during the same condition (41). Our sRNA expression and degradome tag analyses further confirms the previous finding that *C*. *rosea* can adjust its transcriptomic response, and thereby its mycoparasitic interaction mechanisms, to the identity of its mycohost. We further provide strong evidence that sRNAs plays an essential role in this regulation.

The common response against both mycohosts includes enrichment of degradome tags from genes coding for membrane transporters known for their important role in antagonistic interactions (36, 39, 48–50) . The result confirms that the regulation of genes coding for membrane transporters is an important common response of *C*. *rosea* during interspecific fungal interactions (38, 39, 41) and that sRNAs mediate this process. Additionally, enrichment of genes encoding membrane proteins among the sRNA targets points to a drastic replacement of the protein part of the membrane during non-self-interactions of *C. rosea*. The degradome tags of genes putatively associated with biosynthetic processes, including lipid biosynthesis were enriched specifically against *F*. *graminearum.* This suggests that *C. rosea* de-suppresses the expression of lipid biosynthesis genes, possibly to maintain the functionality of the plasma membrane that may be a target for toxic metabolites produced by *F. graminearum*. It is known that *Fusarium* spp. produce toxic compounds for interference competition (41, 42). For instance, the mycotoxin DON is shown to down-regulate the expression of chitinase genes associated with the mycoparasitic attack in *Trichoderma* spp. (51). This fits well with the gene expression profile of genes related to specialised metabolite biosynthesis, including DON. DON-biosynthesis genes were down-regulated when *F. graminearum* grew in contact with a *C. rosea* Δ*dcl2* mutant with diminished biocontrol capabilities, in respect to *F. graminearum* interacting with *C. rosea* WT, suggesting that this mycotoxin is needed to overcome the antagonistic activity of *C. rosea* (41). Similar results were reported previously where expression of genes coding for Kp4 killer toxins were up-regulated in *F. graminearum* during the interaction with the mycoparasitic fungus *T. gamsii* (42).

Another plausible explanation of the differential response is related to the degree of antagonistic ability of the mycohosts. The fungal prey is not passively growing towards the mycoparasite. On the contrary, it can actively launch a counter-attack involving fungal cell wall-degrading enzymes, toxic specialized metabolites and production of reactive oxygen species (52). This is also reflected by the result from the *in vitro* dual culture plate confrontation assay, which revealed a growth rate inhibition of *C*. *rosea* during the interaction with *F. graminearum*, but not with *B. cinerea*. The overgrowth rate of *C*. *rosea* on *B*. *cinerea* mycelia was similar to the growth rate in the non-interaction control, suggesting that *C. rosea* can quickly overcome the counter-attack by *B*. *cinerea*. This is verified by the transcriptome-wide comparative degradome analysis between two interaction stages, which showed how the transcript cleavage pattern 24 h after contact with *B. cinerea* was more similar to the *C. rosea* self-interaction control than to the *B. cinerea* contact stage. This suggests that *C. rosea* quickly overcame *B. cinerea* and that the transcript levels were already going back to normalcy 24 h after contact, while the contact and 24 h after contact stages with *F. graminearum* remained very similar to one another. Co-expression analysis further highlights the mycohost and interaction stage-dependent responses of *C*. *rosea*, with modules 10 and 11 showing an expression in CrBc at after contact more similar to the CrCr control than to CrBc at contact. This also emphasizes that the mycelial contact stage is crucial for non-self-interactions for modulation of the mycoparasitic regulatory network in *C. rosea*. Zapparata et al. (42) also showed extensive communication between the mycoparasite *T. gamsii* and *F. graminearum* resulting in transcriptomic modifications in both fungi, even before physical contact.

Another interesting finding is that the number of down-regulated sRNAs is greater than the number of up-regulated ones, and the number of putative targeted transcripts followed a similar pattern. Mechanistically, this suggests that genes encoding proteins involved in the response of *C. rosea* to mycohosts are constantly suppressed by sRNA-mediated RNAi, when the mycohosts are absent. However, in the presence of mycohosts, *C. rosea* reduces the production of sRNAs, thereby inducing the consequent de-suppression (up-regulation) of the gene targets. Among the targets of sRNAs commonly down-regulated (against both mycohosts), we find transcripts coding for secreted proteins, such as the glycoside hydrolase family 18 chitinase ECH42 (CRV2T00001280_1) known for its role in antagonism (53, 54). Other under-degraded targets that could have a role in the mycoparasitic activity of *C. rosea* include an autophagy protein 5 homolog (CRV2T00000627_1), as the deletion of this gene in *T. reesei* causes sensitivity to nutrient starvation, with abnormal conidiophores and reduced production of conidia (55). Moreover, one of the transcripts cleaved by sRNAs down-regulated in CrFg at contact is predicted to code for a catalase isozyme P (CRV2T00015756_1), suggesting that *C. rosea* could use RNA silencing to regulate its resistance to the high oxidative stress levels that are often present during mycoparasitic interactions (52). Transcripts coding for proteins with regulation activity are also predicted to be cleaved, pointing to the fact that the expression of genes not directly targeted by sRNAs could still be under the indirect control of RNAi, as was observed in other studies (14, 41). Such regulating transcripts include the predicted transcription factors CRV2T00007226_1 and CRV2T00009845_1, the transcription initiation factor IID CRV2T00002865_1 and the mitogen-activated protein kinase CRV2T00004440_1. The number of over-degraded transcripts putatively cleaved by up-regulated sRNAs is much lower, especially 24 h after contact. Still, we find the transcript CRV2T00007976_1 among the putative targets involved in glycogen biosynthesis. Glycogen is an energy storage molecule (56), so it may indicate reduced anabolism in favor of mobilization of the available energy for its antagonistic activity. Furthermore, we identified a gene target coding for a helicase of the SNF2 family (CRV2T00004339_1), targeted by cro-mir-9 up-regulated specifically during contact in CrFg, known for regulating transcription by chromatin remodeling through the deposition of H2A.Z (57).

The role of sRNA in cross-kingdom RNAi is established (19, 20). To investigate sRNA mediated cross-species RNAi in mycoparasitic interaction, mycohost transcripts were also predicted as possible targets (cross-species gene targets) by *C. rosea* sRNAs and milRNAs. Among these, it is exciting to note that *C. rosea* sRNA ii_seq_9854567_x248 (up-regulated at contact and after contact in CrFg) was predicted to target *F. graminearum* transcript XM_011326499.1 coding for a Rab GDP-dissociation inhibitor. The homologue of this gene, whose deletion is lethal, is shown to be involved in chitin deposition in *Aspergillus niger* (58), and chitin degradation is an important part of *C. rosea* antagonistic interactions (31, 53), making the sRNA-based cleavage of transcript XM_011326499.1 an interesting hypothesis (**Table 4**). Similarly, up-regulated milRNA cro-mir-72 was predicted to cleave *F. graminearum* transcript XM_011323146.1, coding for an elongation factor involved in resistance to oxidative stress in *S. cerevisiae* (47). It suggests that *C. rosea* can use RNA silencing to decrease oxidative stress resistance of its mycohosts. The milRNA cro-mir-9, up-regulated during contact with *F. graminearum*, also had two putative cross-regulated targets in Piombo et al. (41), and they encoded a zinc-binding protein (XM_011320613.1) and a sec1 family vesicle transport protein (XM_011329717.1). This family is essential for SNARE-mediated membrane fusion in eukaryotes (59) and is therefore involved in a vast array of biological processes, including virulence in *Cryptococcus neoformans* (60). Moreover, cro-mir-4 showed up-regulation at contact in CrBc, and this milRNA was suggested to have several putative cross-regulated targets in a previous study (41). These include the GT2 chitin synthase BcCHSIV and a homolog of the pp-z protein (XM_024693385.1). This protein is involved in oxidative stress response and virulence in *Candida albicans* (61), and oxidative stress is often present at mycoparasitic interaction sites (52). Another putative target was the vacuolar sorting protein *Bcvps27*, whose homolog deletion in *B. cinerea* causes a reduction in growth rate, aerial hyphae formation and hydrophobicity, as well as increasing sensitivity to cell wall-damaging agents and to osmotic stresses (62). Furthermore, several genes involved in gene expression regulation are also predicted to be targeted by cro-mir-4, suggesting that *C. rosea* can affect *B. cinerea* gene regulation at a higher level to carry out its antagonistic activity. These genes include the chromatin binding factor *Bcyta7* (63) and the transcription factor *Bcltf3*, necessary for conidiogenesis (64) (**Table 3)**. It must be underlined that additional evidence is needed to validate the cross-regulation events.

## Conclusions

The presented work increases our understanding of the mechanisms involved in interspecific fungal interactions, with important implications for the use of fungi as biological control agents. We show that several *C. rosea* sRNAs were down-regulated during interactions with *B. cinerea* and *F. graminearum*. Consequently, their putative gene targets were up-regulated (de-suppressed), suggesting a role of sRNA-mediated regulation of mycoparasitism in *C. rosea.* These putative *C. rosea* sRNA-regulated transcripts were often coding for membrane transporters or secreted proteins, and had a role in resistance to oxidative stress, cellular metabolism and lipid biosynthesis. We further show that the response of *C*. *rosea* towards *B. cinerea* and *F. graminearum* depends on mycohost identity and interaction stage, and that sRNAs are part of the regulatory mechanism. This is important as it shows that *C. rosea* can adapt its transcriptional response, and thereby its interaction mechanisms, based on the identity of the mycohost. Finally, our data strongly suggest a role of cross-species RNAi in mycoparasitism, representing a novel mechanism in biocontrol interactions. This can find important applied uses in spray-induced gene silencing for improved efficacy of biocontrol applications.

## Material and methods

### Experimental setup for small RNA and degradome sequencing

*Clonostachys rosea* strain IK726, *B. cinerea* strain B05.10 strain and *F. graminearum* strain PH-1 were used in the study. An *in vitro* dual culture experimental was performed for sRNA and degradome sequencing during the interaction, following previously described procedures (35). In brief, an agar plug of *C*. *rosea* mycelium was inoculated at the edge of a 9-cm-diameter potato dextrose agar (PDA; Merck, Kenilworth, NJ) Petri plate covered with a durapore membrane filter (Merck, Kenilworth, NJ) for an easy harvest of mycelia. The mycohosts fungi *B. cinerea* or *F*. *graminearum* were inoculated at opposite sides of the plate (37). The mycelial front (5 mm) of *C*. *rosea* was harvested together with the mycelial front (5 mm) of *B*. *cinerea* (CrBc) or *F*. *graminearum* (CrFg) at the hyphal contact stage (early physical contact between the mycelia), and at 24-hour post hyphal contact stage of interactions. Mycelium harvested at the same stage from *C*. *rosea* confronted with *C*. *rosea* (CrCr) was used as a control treatment. The experiment was performed in four biological replicates.

### RNA extraction and sequencing

Total extraction was performed using the mirVana miRNA isolation kit following the manufacturer’s protocol (Invitrogen, Waltham, MA). The RNA quality was analysed using a 2100 Bioanalyzer Instrument (Agilent Technologies, Santa Clara, CA) and RNA concentration was quantified using a Qubit fluorometer (Life Technologies, Carlsbad, CA). For sRNA sequencing, the total RNA was sent for library preparation and paired-end sRNA sequencing at the National Genomics Infrastructure (NGI) Stockholm. The sRNA library was generated using TruSeq small RNA kit (Illumina, San Diego, CA) and sequenced on one NovaSeq SP flowcell with 2×50 bp reads using Illumina NovaSeq6000 equipment at NGI Stockholm. The Bcl to Fastq conversion was performed using bcl2fastq_v2.19.1.403 from the CASAVA software suite. The quality scale used is Sanger / phred33 / Illumina 1.8+.

### Functional annotation of genomes

BLAST2GO v.5.2.5 (65) and InterProScan v.5.46-81.0 (66) were used to annotate the proteomes of *C. rosea* strain IK726 (PRJEB4200), *B. cinerea* strain B05.10 (ASM14353v4) and *F. graminearum* strain PH-1 (ASM24013v3). Putative CAZymes were identified through the dbCAN2 meta server (67).

### Small RNA sequence analysis

All the analysis done on sRNAs is summarize in **Figure 9**. Raw reads received after sequencing were trimmed with cutadapt v. 2.8 (68), setting 16 bp as the minimum allowed length, and quality was checked with FastQC v.0.11.9 (https://www.bioinformatics.babraham.ac.uk/projects/fastqc/) before and after the trimming. Reads shorter than 18 bp or longer than 32bp were removed, and the cleaned reads were then mapped to the genomes of *C*. *rosea* strain IK726 (69), *B*. *cinerea* strain B05.10 (70) and *F*. *graminearum* strain PH-1 (71) using bowtie v.1.2.3 (72) with the following parameters: bowtie -p 30 -n 1 -l 16 -m 200 --best --strata --chunkmbs 500. To exclude the sRNA sequences originating from the mycohosts, the reads mapping exclusively to the *C*. *rosea* genome were retained and were used for further analysis. Sense and antisense sRNAs mapping to intergenic regions and introns, as well as antisense sRNAs mapping to exons, were detected and DESeq2 v.1.28.1 (73) was used for differential sRNA expression analysis at *P*-value 0.05 at the cutoff of log2 fold change (FC):1 and *P*-value (adj) = 0.05. Moreover, milRNAs were predicted with miRDeep2 with default parameters (46). Both known and novel milRNAs were predicted, and DESeq2 v.1.28.1 (73) was used to determine differential expression.

**Figure 9.**
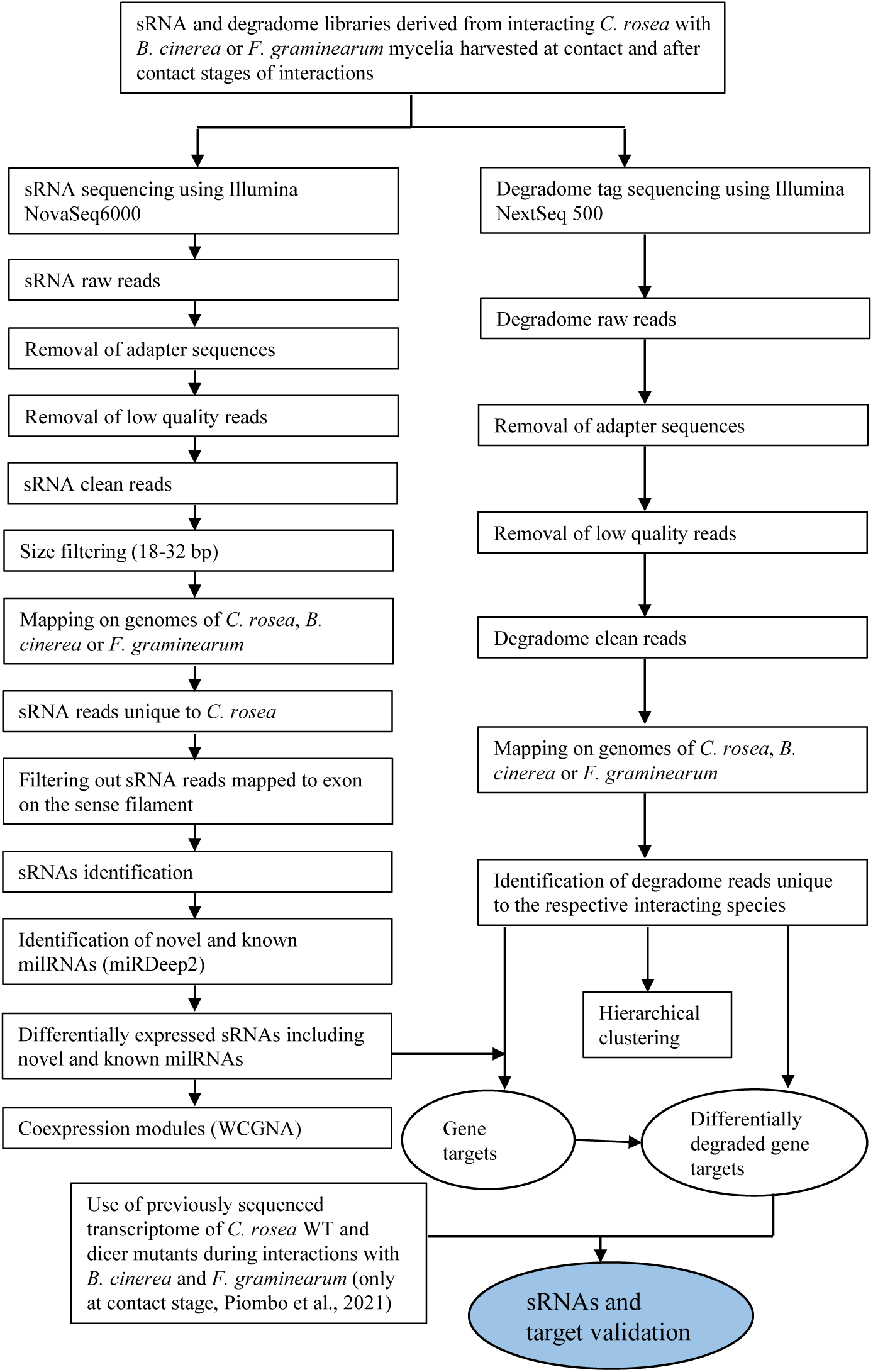
The flowchart of sRNAs and degradome sequence analyses showing different steps in identifying *C. rosea* sRNAs (including milRNAs) and their potential endogenous and cross-species gene targets.

To ensure the novelty of newly detected milRNAs, they were compared with the fungal milRNAs identified in several other studies, plus all the fungal milRNAs available in RNAcentral, using the Blast algorithm with 95% minimum identity (13, 16, 17, 74–78).

### Coexpression analysis

Normalised expression values were obtained for sRNAs by using DESeq2 (73). The values of differentially expressed sRNAs were used to perform a coexpression analysis with WCGNA (79) using a soft-thresholding power of 6. The function “binarizeCategoricalVariable” was used to convert the mycohost and interaction stage categorical variables into numerical ones, and spearman correlation was calculated between them and the module eigengenes.

### MilRNA target prediction

The UTR regions of *B. cinerea*, *F. graminearum* and *C. rosea* genes were determined with “add_utrs_to_gff” (https://github.com/dpryan79/Answers/tree/master/bioinfoSE_3181), and they were used for target prediction analysis for animal-based tools PITA, Miranda, TargetSpy and simple seed analysis within the sRNA toolbox (80). The sRNA toolbox was also used to run the plant-based target predictors psRobot and TAPIR (81, 82), while Targetfinder and psRNATarget were used independently (83, 84) . Target-milRNA couples predicted by at least 3 animal-based tools or 2 plant-based ones were considered in the following analyses.

### Degradome sequencing and analysis

To sequence uncapped 5’ ends from poly-adenylated RNA (Degradome Seq), total RNA isolated from the abovementioned samples was used. For degradome sequencing, total RNAs, after DNAse treatment, was sent to GenXPro (GenXPro GmbH, Frankfurt, Germany). The degradome libraries were generated by GenXPro using the MACE-Seq Kit (Massive analysis of cDNA ends) (GenXPro GmbH, Frankfurt, Germany) and the TrueQuant small RNA-Seq Kit (GenXPro GmbH, Frankfurt, Germany). Briefly, mRNA was captured using Dynabeads OligodT (Invitrogen, Waltham, MA). The 5’ adapter from the TrueQuant small RNA-Seq kit (GenXPro GmbH, Frankfurt, Germany) was ligated to the uncapped 5’ ends of the poly-A transcripts. Reverse transcription and PCR was performed according to the TrueQuant small RNA-Seq kit manual, and the degradome Tags were sequenced on an Illumina Next 500 insteument. Reads were trimmed with bbduk v.38.86 (85) with the following options: bbduk.sh in1=read1.fastq in2=read2.fastq out1=read1_clean.fastq out2=read2_clean.fastq ref=.fa ktrim=r k=23 mink=11 hdist=1 tpe tbo qtrim=r trimq=10. The analysis of degradome data is summarized in **Figure 9**.

Successful cleaning and adapter removal were verified with FastQC v. 0.11.9 (https://www.bioinformatics.babraham.ac.uk/projects/fastqc/). Since all the samples represented the interaction of two organisms, the genome of *C. rosea* was concatenated with the one of either *B. cinerea* or *F. graminearum*, creating two combined genome files (CrFg and CrBc), and the same was done with the annotations in gff format. Degradome reads from *C. rosea*-*B. cinerea* interaction were aligned to the CrBc genome, while reads from *C. rosea*-*F. graminearum* interactions were aligned to the CrFg one. Multimapping reads were removed from the analysis. The chosen aligner was STAR v.2.7.5c (86), with default options, and the count tables were then generated through featureCounts v.2.0.1 (87). Only sense reads were considered for mapping to known features, while both sense and antisense reads were considered when mapping to intergenic regions. Variance stabilising transformation was applied to visualise the hierarchical clustering with the R pheatmap package (88).

Differentially expressed sRNAs mapping to intron or intergenic regions, and antisense sRNAs mapping to exons, were used together with degradome reads to predict which genes were regulated through RNA silencing, by using the Cleaveland program v. 4.5 (11). Only genes flagged as category two or better degradation targets in all replicates were retained as putative targets of RNA silencing. Category 2 means that a higher than average degradome read count was present at the mapping site of the considered sRNA (https://github.com/MikeAxtell/CleaveLand4). After this step, we estimated differential degradation at the point of alignment between each sRNA and its target, using DESeq2 with default parameters and as input only counts of degradome reads mapping to the point of alignment predicted by Cleaveland. We retained only the results in which the differential degradation and the differential sRNA expression showed correlation (for example underdegradation in targets of down-regulated sRNAs). Furthermore, the expression level of transcripts putatively cleaved by sRNAs was checked using the data from Piombo et al. (41), verifying how many of the putative targets were up-regulated in *C. rosea* Dicer deletion mutants, devoid of a functional Dicer-dependent RNA silencing system.

Enrichment in GO terms in the set of genes targeted by differentially expressed sRNAs was determined through Fisher tests performed with AGRIGO (89) using the Yekutieli Multi-test adjustment method and an FDR threshold of 0.05. The results were then visualized through REVIGO (90).

### PhasiRNA prediction

PhasiRNAs in the dataset were predicted with PHASIS v. 3 (91) setting minimal abundance to 10, and differential expression was analysed with DESeq2 v.3.13 (73). Target prediction was carried out with Targetfinder, psRobot, TAPIR and psRNAtarget, and only targets co-predicted by at least 2 tools were considered (80–84).

### milRNAs detection in other *Clonostachys* spp.

The presence of novel and known milRNAs were investigated in the genomes of *C. byssicola* CBS 245.78, *C. chloroleuca* CBS 570.77, *Clonostachys* sp. CBS 192.96, *C. rhizophaga* CBS 906.72A and *C. solani* 1703 (48). The analysis was done through Blast with the option “-- task blastn-short”, using a 95% threshold in both identity and query coverge for milRNA mature sequences, and 90% for the entire precursor sequences.

### Stem loop RT-qPCR

Stem loop RT-qPCR primers were designed for specific miRNAs (92) (supplemental table S11. Prior to reverse transcription, the stem loop primers were denatured by incubating at 65 °C for 5 minutes and transferring to ice immediately. Reverse transcription was carried out by adding the denatured stem loop primer (final conc. 0.05 µM) to the following components – dNTPs (final conc. 0.25 mM), 10× SS III buffer (final conc. 1×), dithiothreitol or DTT (final conc. 10 mM), RNaseOUT RNA inhibitor (final conc. 0.2 U/µL), Superscript III reverse transcriptase enzyme (final conc. 2.5 U/µL, Invitrogen, Waltham, MA), reverse primer for *C. rosea* reference gene Actin (final conc. 0.25 µM). RNA (10 ng) from respective control and treatment samples were used as template and the reaction volume was made up to 20 µL using nuclease-free water. In a thermal cycler, the following reaction conditions were used – 16 °C incubation for 30 minutes; 60 cycles consisting of 30 °C for 30 seconds, 42 °C for 30 seconds and 50 °C for 1 second. The reaction was then terminated by enzyme inactivation at 85 °C for 5 minutes.

RT-qPCR was performed with the help of DyNAmo Flash SYBR green kit (Thermo Fisher Scientific, Waltham, MA) to validate the relative miRNA expression. *C. rosea* Actin was included as a reference gene for normalization. The Livak method (2^-ΔΔCt^) was employed for quantification of gene expression (93).

## Acknowledgements

This work was financially supported by the Department of Forest Mycology and Plant Pathology; Swedish Research Council for Environment, Agricultural Sciences and Spatial Planning (FORMAS; grant number 2018-01420), and Carl Tryggers Stiftelse för Vetenskaplig Forskning (CTS 19: 82). MK acknowledges SLU Centre for Biological Control (CBC) at the Swedish University of Agricultural Sciences. RV is supported by FORMAS (2019-01316), Carl Tryggers Stiftelse för Vetenskaplig Forskning (CTS 20: 464), Partnerskap Alnarp and the Crafoord foundation (20200818). The authors acknowledge support from the National Genomics Infrastructure in Stockholm funded by Science for Life Laboratory, the Knut and Alice Wallenberg Foundation and the Swedish Research Council, and SNIC/Uppsala Multidisciplinary Center for Advanced Computational Science for assistance with massively parallel sequencing and access to the UPPMAX computational infrastructure.

## Supplementary materials

**Supplementary table S1**: Summary of sRNA sequencing

**Supplementary table S2:** Differentially expressed sRNAs in *Clonostachys rosea* during interaction with *B. cinerea* (CrBc) and *F. graminearum* (CrFg) compared with CrCr control, at contact (C) and after contact (AC) stages of interactions.

**Supplementary table S3:** Sequence, abundance, origin and genome coordinates of the milRNAs detected in this study.

**Supplementary table S4**: Summary of degradome sequencing and percentage of degradome reads mapping to genome features.

**Supplementary table S5:** Endogenous transcripts (endogenous gene targets) putatively cleaved by differentially expressed sRNAs.

**Supplementary table S6:** Putative cross-species gene targets in *B. cinerea* and *F. graminearum* putatively cleaved by differentially expressed *C. rosea* sRNAs.

**Supplementary table S7**: Putative endogenous gene targets of differentially expressed milRNAs.

**Supplementary table S8**: Putative cross-species gene targets in *F. graminearum* putatively cleaved by differentially expressed *C. rosea* milRNAs.

**Supplementary table S9**: Distribution of the precursor and mature sequences of *C. rosea* novel and known milRNAs in *Clonostachys* spp.

**Supplementary table S10**: Sequence, origin and coordinates of the phasiRNA families detected in this study.

**Supplementary table S11**: List of primers used in stem-loop RT-qPCR.

**Supplemental figure S1:** Schematic illustration of *in vitro*-dual culture plate confrontation assay used in this study for sRNAs and degradome sequencing experiment. (**A**) contact stage, (**B**) after contact stage of interactions.

**Supplementary figure S2**: Growth rate of *C. rosea*, *B. cinerea* and *F. graminearum* on PDA. **(A**) Growth rate of *C. rosea* during non-interaction control (Cr), self-interaction (CrCr) and non-self-interaction with *B. cinerea* (CrBc) and *F. graminearum* (CrFg). (**B**) The growth rate of *B. cinerea* during non-interaction control (Bc), self-interaction (BcBc) and non-self-interaction with *C. rosea* (CrBc). (**C**) The growth rate of *F. graminearum* during non-interaction control (Fg), self-interaction (FgFg) and non-self-interaction with *C. rosea* (CrFg). Agar plugs were inoculated on opposite sides in 9 cm diameter agar plates and incubated at 25°C. Growth rates of *C. rosea*, *B. cinerea* and *F. graminearum* were recorded daily two days post inoculation (dpi) until the mycelial contact. Due to the slower growth rate of *C. rosea*, *B. cinerea*, or *F. graminearum* were inoculated 7dpi in non-self-interaction experiments.

**Supplementary figure S3**: Validation degradome based gene targets by transcriptome sequencing. Percentage of genes, putatively cleaved by differentially expressed sRNAs in this study, were also up-regulated in the Δ*dcl2* mutant during interaction with *B. cinerea* and *F. graminearum* in Piombo et al (41). There was no degradome-based gene target to validate up-regulated sRNA at afar contact stage of CrBc (CrBc_AC_up-regulated).

